# Cdk9 regulates a promoter-proximal checkpoint to modulate RNA Polymerase II elongation rate

**DOI:** 10.1101/190512

**Authors:** Gregory T. Booth, Pabitra K. Parua, Miriam Sansó, Robert P. Fisher, John T. Lis

**Affiliations:** Department of Molecular Biology and Genetics, 107 Biotechnology Building, 526 Campus Road, Cornell University, Ithaca, NY 14853-2703, USA; Department of Oncological Sciences, Icahn School of Medicine at Mount Sinai, New York, NY 10029, USA

## Abstract

Multiple kinases modify RNA Polymerase II (Pol II) and its associated pausing and elongation factors to regulate Pol II transcription and transcription-coupled mRNA processing^1,2^. The conserved Cdk9 kinase is essential for regulated eukaryotic transcription^3^, but its mechanistic role remains incompletely understood. Here, we use altered-specificity kinase mutations and highly-specific inhibitors in fission yeast, *Schizosaccharomyces pombe* to examine the role of Cdk9, and related Cdk7 and Cdk12 kinases, on transcription at base-pair resolution using Precision Run-On sequencing (PRO-seq). Within a minute, Cdk9 inhibition causes a dramatic reduction in the phosphorylation of Pol II-associated factor, Spt5. The effects of Cdk9 inhibition on transcription are the more severe than inhibition of Cdk7 and Cdk12 and result in a shift of Pol II towards the transcription start site (TSS). A kinetic time course of Cdk9 inhibition reveals that early transcribing Pol II is the most compromised, with a measured rate of only ~400 bp/min, while Pol II that is already well into the gene continues rapidly to the end of genes with a rate > 1 kb/min. Our results indicate that while Pol II in *S. pombe* can escape promoter-proximal pausing in the absence of Cdk9 activity, it is impaired in elongation, suggesting the existence of a conserved global regulatory checkpoint that requires Cdk9 kinase activity.

---

Recently, we reported promoter-proximal pause-like distributions of Pol II on many genes in fission yeast^4^, yet whether such pausing is regulated through kinase activity is not known. In metazoans, kinase inhibitors, including 5,6-dichloro-1-β-D-ribofuranosylbenzimidazole (DRB) and flavopiridol (FP), which primarily inhibit Cdk9, have been instrumental in the discovery of promoter-proximal pausing as a major regulatory hurdle for most genes^5-8^. However, off-target effects of chemical inhibitors make selective ablation of Cdk9 activity *in vivo* nearly impossible^9^.

To isolate the immediate impact of loss of Cdk9 activity on Pol II pausing and elongation at single nucleotide resolution in fission yeast, we performed PRO-seq in an analog sensitive mutant (AS) strain, *cdk9*^*as*^, which is vulnerable to inhibition by the addition of 3-MB-PP1^10^. Phosphorylation of Spt5 was undetectable after one minute of drug addition in *cdk9*^*as*^ (Fig. 1a), while CTD residues were largely unaffected, within the timeframe tested (Extended Data Fig. 1a). The near-immediate loss of Spt5 phosphorylation upon Cdk9 inhibition supports a model of active, competing Spt5 dephosphorylation (see accompanying paper, Parua *et. al.* 2017), and a critical influence of this mark on transcription. PRO-seq libraries were prepared in biological replicates (Extended Data Fig. 1b) from *cdk9*^*as*^ cells treated for 5 minutes with 10 μM 3-MB-PP1 (treated) or an equivalent volume of DMSO (untreated). As a control we prepared PRO-seq libraries from equivalently treated wild-type (*wt*) cells. Comparison of untreated *cdk9*^*as*^ profiles with *wt* profiles revealed no obvious differences (Extended Data Fig. 2a) and addition of 3-MB-PP1 to *wt* cells produced almost no transcriptional changes (Extended Data Fig. 2b), indicating that this system has little to no basal phenotypes of the AS mutation or off-target drug effects. To evaluate the effect of Cdk9 on promoter-proximal pausing, genes were divided into quartiles based on their pausing index (PI), which measures the enrichment of Pol II in the promoter-proximal region (TSS to +100 nt) relative to the gene body. Although we observed striking changes in average gene body PRO-seq signal resulting from 5 minutes of *cdk9*^*as*^ inhibition, pausing did not explain this effect (Fig. 1b).

**Figure 1.**
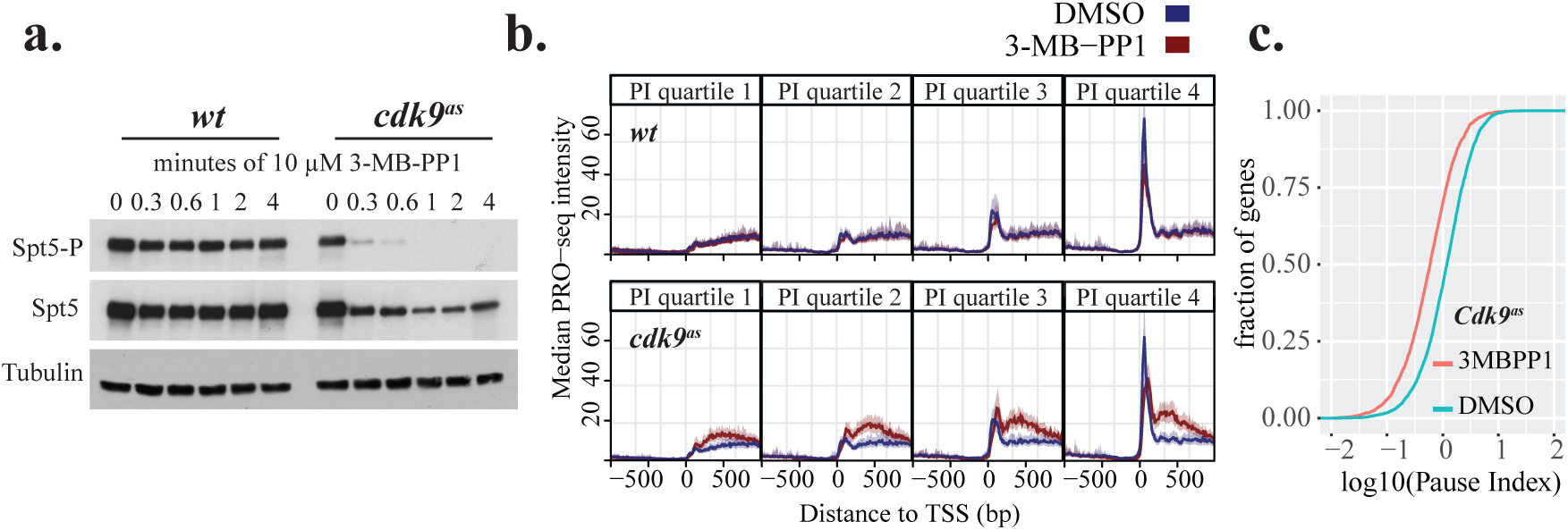
Loss of Cdk9 activity does not prevent escape from promoter-proximal pause during transcription elongation. **a**. Western blot analysis of pSpt5 compared with total Spt5 in *wt* and *cak9*^*as*^ over a time course of cultures treated with 10 μ*M* 3-MB-PP1. Tubulin serves as a loading control. **b**. TSS-centered composite profiles of PRO-seq data before (blue) and after (red) 5 minutes of treatment with 10 μM 3-MB-PP1 for genes grouped by quartiles of increasing pausing index from left to right (calculated using untreated wild type data from combined replicates). The top four panels represent wild-type data, while the bottom four panels represent *cdk9*^*as*^ data. **c.** Cumulative density functions for pausing index (log_10_) of all filtered genes in treated (red) and untreated (blue) samples for the *cak9*^*as*^ strain.

Inhibition of pause escape in mammals is known to result in an increase in engaged, promoter-proximal Pol II coinciding with loss of signal from the gene body^6^, yielding a greater PI for many genes. In contrast, we find few genes with significant increases in promoter proximal Pol II upon Cdk9 inhibition (Extended Data Fig. 2c & d). Moreover, compared to *wt*, *cdk9*^*as*^ cells exhibit a global decrease in pausing index as a result of 5 minutes of treatment with 3-MB-PP1 (Fig. 1c, Extended Data Fig. 2e). Together, our results reveal that Cdk9 activity in *S. pombe* is not critical for release of paused Pol II into elongation, but it is nonetheless critical for normal transcription.

CTD modification by transcription-coupled kinases, Mcs6 (Cdk7) and Lsk1 (Cdk12)^11^ can modulate the association of auxiliary components with Pol II, facilitating co-transcriptional RNA processing^2^ and elongation through chromatin^12^. Thus, in addition to Cdk9, we set out to measure the impact of Mcs6 and Lsk1 on global transcription. A novel AS variant of Mcs6 (here called *mcs6*^*as5*^) was as sensitive to treatment with 3-MB-PP1 as *cdk9*^*as*^ (Extended Data Fig. 3a, b). Importantly, no gross transcriptional differences were observed in untreated *mcs6*^*as5*^ compared with *wt* (Extended Data Fig. 2a). Within minutes of treatment, we observed measurable losses in Ser5-P and Ser7-P in the *mcs6*^*as5*^ strain, while these marks in *wt* cells were unaffected (Fig. 2a). The analog sensitive Lsk1 (*lsk1*^*as*^) was also inhibited by 10 μM 3-MB-PP1 (Extended Data Fig. 3c). However, despite the clear impact on Ser2 phosphorylation (Extended Data Fig. 4a), there was almost no visible change in transcription upon treatment of *lsk1*^*as*^ with 10 μM 3-MB-PP1 for 5 minutes (Extended Data Fig. 4b-d). In contrast, loss of Mcs6 or Cdk9 activity produced abrupt changes at both ends of individual genes (Fig. 2b). Most notably in *cdk9*^*as*^, Pol II signal tapered off towards the 3’ end of the transcription unit after inhibition. Moreover, combined inhibition of either *lsk1*^*as*^ and *cdk9*^*as*^, or *mcs6*^*as5*^ and *cdk9*^*as*^ produced transcriptional defects closely resembling those of *cdk9*^*as*^ alone, suggesting an overriding importance of Cdk9 in maintaining normal elongation in *S. pombe* (Extended Data Fig. 5).

**Figure 2.**
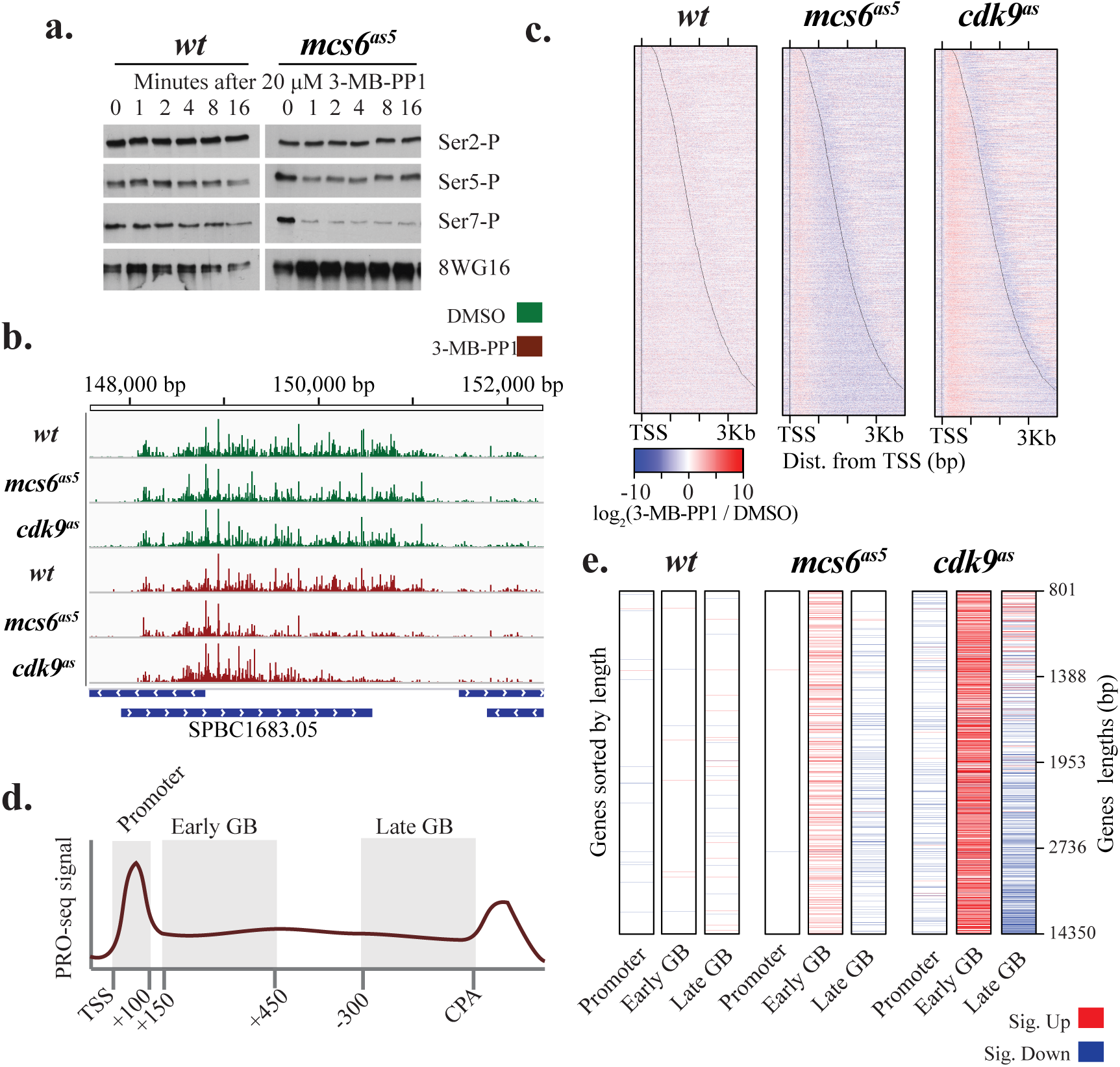
Cdk9 inhibition has a more severe impact on global transcription elongation than Mcs6 inhibition. **a**. Western blot analysis of phosphorylated CTD residues (Ser2-P, Ser-5-P, Ser7-P) in relation to total CTD signal. Levels were measured over a time course of cultures treated with 20 μM 3-MB-PP1. **b**. Browser track image from the *SPBC1683.05* locus displaying normalized read counts from untreated (green) and treated data (red) for wild-type, *mcs6*^*as5*^ and *cdk9*^*as*^ strains from top to bottom, respectively (plus strand only). All PRO-seq samples represent treatments for 5 minutes with 10 μM 3-MB-PP1 (treated) or equivalent volume of DMSO (untreated). **c**. Heat maps depicting the log_2_ fold change in normalized PRO-seq signal (treated/untreated) within 10 bp bins from -250 bp to +4000 bp relative to the TSS for wild-type, *mcs6*^*as*^ and *cdk9*^*as*^ strains from left to right, respectively. Genes within each heat map are sorted by increasing length from top to bottom, with black lines representing observed TSS and CPS. **d**. Illustration of the definition of promoter (TSS to +100) early (+150 to +450 from TSS) and late (-300 to 0 from CPS) gene body regions used in E. **e**. Heat maps depicting whether each gene exhibits a significant fold change (adjusted *p* < 0.01; treated/untreated) in promoter, early or late gene body regions. Genes were required to be longer than 800 nt and are sorted from top to bottom by increasing gene length with length quartiles shown on the right. Significant increases and decreases in each region are shown as red and blue, respectively.

Composite PRO-seq profiles around transcription start sites (TSS) and cleavage and polyadenylation sites (CPS) revealed global kinase-dependent effects at both ends of transcription units (Extended Data Fig. 5). Surprisingly, the loss of signal at gene 3’ ends upon inhibition of Mcs6 or Cdk9 was only observed on longer genes (Fig. 2c). Differential expression analysis ^13^ was used to verify differences in treated and untreated PRO-seq counts within 3 discrete regions of each gene: 1) the promoter (TSS to +100), 2) the early gene body (+150 to +450 from the TSS), and 3) the late gene body (-300 to CPS) (Fig. 2d). Most strikingly in *cdk9*^*as*^, and to some extent in *mcs6*^*as5*^, when genes were sorted by increasing length, distance from the TSS appeared to dictate the pattern of changes within the gene body regions. Although promoter-proximal regions were only subtly affected, early gene bodies were consistently found to show increases in signal in response to Cdk9 or Mcs6 inhibition. However, late gene bodies displayed results relating to the region’s distance from TSS, where short genes exhibited increased signal and long genes had decreased signal within the last 300 bp of the transcription unit (Fig. 2e).

Several mechanisms might explain the observed gene length-dependent effects on transcription. In the absence of Cdk9, Pol II may exhibit a compromised processivity^14^ preventing transcription elongation beyond a certain distance from the TSS. Alternatively, the altered transcription profiles may reflect a reduced rate of early transcribing ECs incapable of, or delayed in, accelerating to the natural productive speed.

We reasoned that these proposed explanations could be resolved by examining a time course of Pol II distribution on genes following rapid Cdk9 inactivation. For instance, a defect in Pol II processivity would result in premature termination at roughly the same distance from the TSS, regardless of time after 3-MB-PP1 addition, whereas a rate defect might reveal a downstream-shifting “wave front” with time of inhibition. Thus, we performed a high-resolution time course of treatment with 3-MB-PP1 followed by PRO-seq, focusing specifically on the severe phenotype in *cdk9*^*as*^.

Strikingly, individual genes reproducibly exhibited a shifting distribution of transcribing Pol II, which appeared to propagate from the TSS toward the 3’ ends of genes over time (Fig. 3a, Extended Data Fig. 6a-c). Heat maps of fold change after each duration of 3-MB-PP1 exposure, relative to untreated (DMSO), revealed an immediate global impact of Cdk9 inhibition on transcription that progressed as a spreading of increased signal across genes over time. Meanwhile, an early drop in signal at gene 3’ ends was gradually recovered with the advancing Pol II density (Fig. 3b).

**Figure 3.**
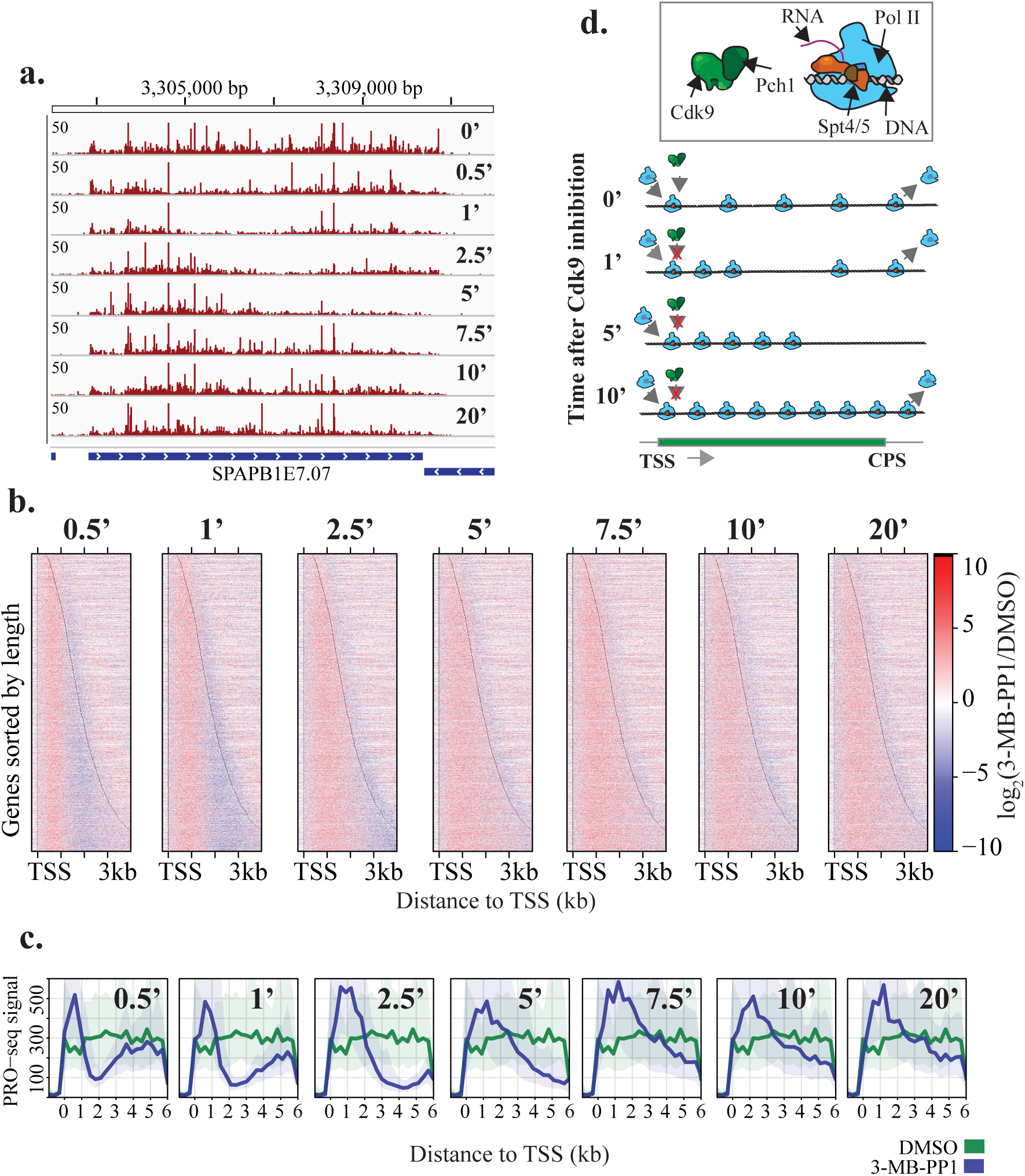
Cdk9 impacts transcription rates through a step during early elongation. **a**. Browser track image displaying the normalized PRO-seq read count signal at the *SPBAPB1E7.07* locus for *cdk9*^*as*^ cells treated with 10 uM 3-MB-PP1 for an increasing duration, from top to bottom. **b**. Heat maps of log_2_(treated/untreated) normalized PRO-seq signal within10 bp windows from -250 bp to +4000 bp around the TSS for all filtered genes ordered by increasing gene length from top to bottom. Panels show PRO-seq data from each time point of drug treatment (relative to the 0 minute sample), with increasing duration from left to right. **c**. Composite PRO-seq signal for all filtered genes at least 6 kb in length (n= 38) before and after treatment. Panels from left to right show profiles after increasing duration of treatment compared with DMSO treated cells. **d**. Illustration of two populations of transcribing Pol II, which have rates differentially affected by Cdk9 inhibition.

To better understand the propagation of increased Pol II density following the initial loss in downstream signal, we prepared composite profiles at each time-point considering only the longest genes in our filtered set (at least 6 kb, n = 38). Intriguingly, average profiles for these genes reveal a fleeting recession of Pol II density towards the CPS (Fig. 3c) - a phenomenon also obvious within individual genes (Fig. 3a; see 1-minute and 2.5-minute time-points). This rapid clearing of gene bodies is reminiscent of the affects of FP in mammals, where late transcribing polymerases continue rapidly to clear off genes, apparently unaffected by the drug^6^. Although not fixed at a specific pause position as in mammals^15^, early transcribing complexes (< 1kb from the TSS) appear to proceed with a decreased elongation rate after the inhibition of Cdk9, indicative of a critical regulatory step during the early stages of elongation in fission yeast (Fig. 3d). Additionally, our transient detection of an unaffected population, rapidly elongating beyond the CPS, suggests differences in transcription rate, rather than premature termination, are the cause of downstream clearing.

Gene activation and repression have been used to derive transcription kinetics in mammals by following changes in Pol II distributions over time^6,16,17^. By adapting a previously described model^18^, our high temporal and spatial resolution data allowed us to determine the distance covered by the advancing waves accurately, despite the comparatively short lengths of *S. pombe* genes (Fig. 4a). For 43 filtered genes longer than 4 kb we were able to estimate advancing wave distance at every time point from 30 seconds to five minutes based on the differences between treated and untreated signals within tiled windows across each gene (Extended Data Fig. 7a). Consistent with a forward moving population of Pol II, distance estimates increased with time of Cdk9 inhibition (Fig. 4b). Moreover, by fitting a linear regression to distance travelled over time elapsed, we determined that the average rate of transcription in this population was 376 bp/minute (Fig. 4c, Extended Data Fig. 8a).

**Figure 4.**
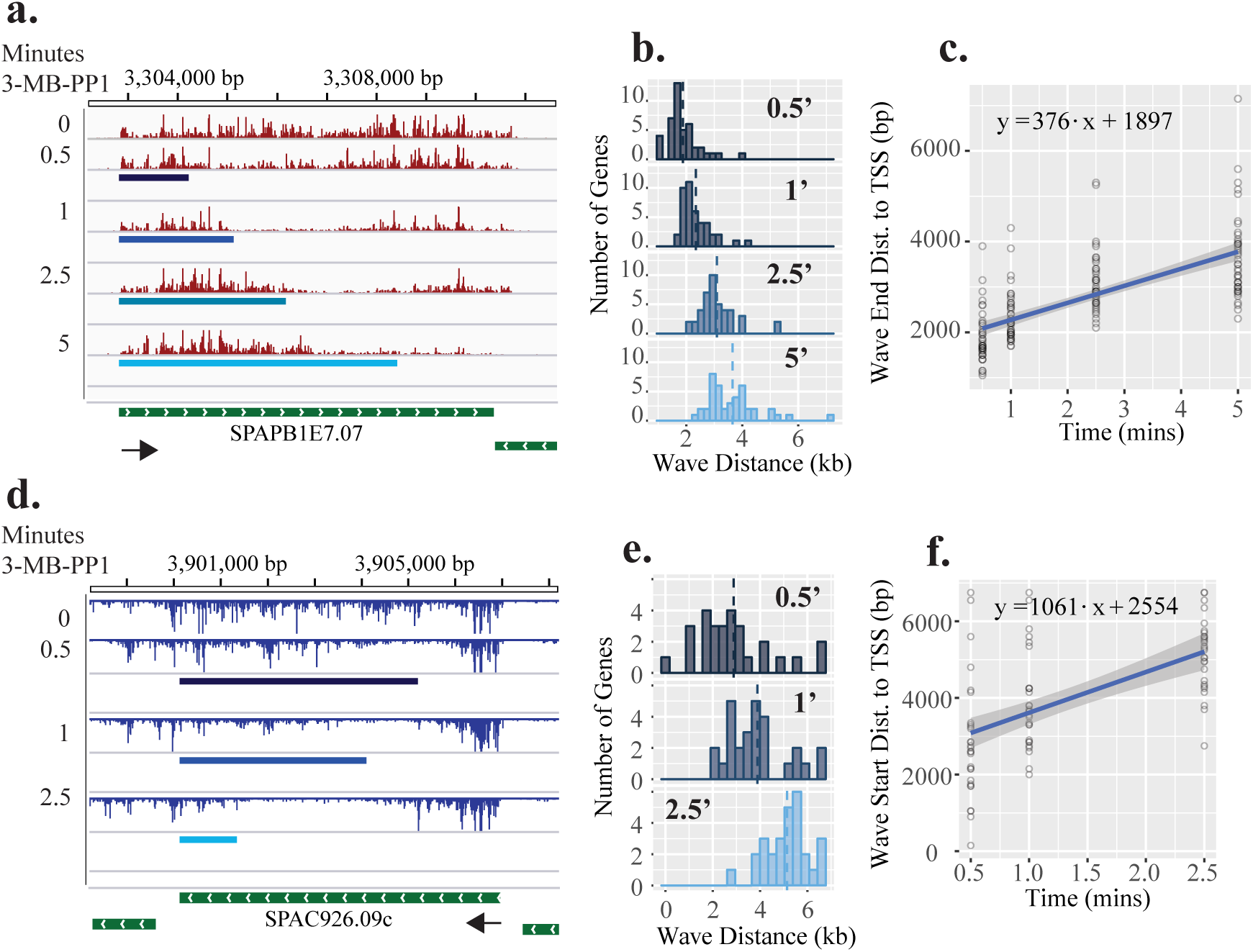
Transcription rate measurments reveal two distinct populations of transcribing Pol II after *cdk9*^*as*^ inhibition. **a**. Browser image displaying normalized PRO-seq signal for increasing lengths of *cdk9*^*as*^ inhibition (top to bottom) over the *SPBAPB1E7.07* locus with advancing wave estimates (blue rectangles) for each time point (0.5 min, 1 min, 2.5 min, 5 min) plotted below the corresponding data. **b**. Histograms of estimated advancing wave end distances (relative to TSS) for filtered genes (n = 43) at each time point after addition of 10 μM 3-MB-PP1 to *cdk9*^*as*^ cells. Dotted, vertical lines represent the mean distance for each distribution. **c**. Linear regression of distances travelled against time elapsed gives the estimated rate of advancing wave progression (slope). **d**. Browser image showing clearing wave measurements (blue rectangles) for each time point plotted below the corresponding data at the *SPAC926.09c* locus. **e**. Histograms of estimated clearing wave tail end distances (relative to CPS) for filtered genes (n = 28) at time points (0.5 min, 1 min, 2.5 min) after *cdk9*^*as*^ inhibition. **f**. Linear regression of clearing wave distances travelled versus time elapsed gives the estimated rate of clearing wave progression (slope).

The observation of a temporarily expanding region of cleared Pol II on long genes suggests that Pol II further into the gene body is moving faster than the advancing wave upon loss of Cdk9 activity. To determine the rate of this “clearing” population of Pol II we further adapted the Danko *et al*. model^18^ to identify the region between the advancing and clearing waves, which thus contains the start of the clearing wave (Extended Data Fig. 7b). Because the clearing population is fleeting and only present on the longest genes, we were forced to omit the five-minute time point and further restrict our analysis to 28 genes longer than 6 kb. Again, clearing wave positions predicted by the model agreed well with PRO-seq profiles (Fig. 4d) and receded away from the TSS on average (Fig. 4e). Moreover, the average transcription rate of clearing complexes was estimated to be 1061 bp/min (Fig. 4f), significantly faster than the advancing wave (Extended Data Fig. 8a). Moreover, analysis of post-CPS elongation after 20 minutes of Cdk9 inhibition shows a narrowed zone of termination (Extended Data Fig. 8b, c, d), consistent with slower moving Pol II being an easier target for exonuclease-mediated termination^19^.

The observation of a temporarily expanding region of cleared Pol II on long genes suggests that Pol II further into the gene body is moving faster than the advancing wave upon loss of Cdk9 activity. To determine the rate of this “clearing” population of Pol II we further adapted the Danko *et al*. model^18^ to identify the region between the advancing and clearing waves, which thus contains the start of the clearing wave (Extended Data Fig. 7b). Because the clearing population is fleeting and only present on the longest genes, we were forced to omit the five-minute time point and further restrict our analysis to 28 genes longer than 6 kb. Again, clearing wave positions predicted by the model agreed well with PRO-seq profiles (Fig. 4d) and receded away from the TSS on average (Fig. 4e). Moreover, the average transcription rate of clearing complexes was estimated to be 1061 bp/min (Fig. 4f), significantly faster than the advancing wave (Extended Data Fig. 8a). Moreover, analysis of post-CPS elongation after 20 minutes of Cdk9 inhibition shows a narrowed zone of termination (Extended Data Fig. 8b, c, d), consistent with slower moving Pol II being an easier target for exonuclease-mediated termination^19^.

Collectively, our results communicate novel mechanistic insight into the importance of co-transcriptional modification of the EC in eukaryotes: 1) Cdk9 plays a dominant role in elongation control, likely reflecting phosphorylation of Spt5 during an early elongation checkpoint^20^, that may be conserved in metazoans, 2) Similarities of the transcriptional response to kinase inhibition suggest other CDKs, particularly Cdk7, might contribute to this regulation^21^, and 3) Cdk9 activity is not essential for pause escape, but crucial for acceleration of early transcribing Pol II. Accordingly, a metazoan-specific chromatin structure or factor, such as the negative elongation factor (NELF) ^22^, which is absent in fission yeast^23^, might further stabilize paused Pol II to create opportunities for precise gene regulation.

## Acknowledgements

Research reported in this publication was supported by NIH grant GM104291 to R.P.F. and GM25232 to J.T.L. The content is solely the responsibility of the authors and does not necessarily represent the official views of the National Institutes of Health. We thank Cornell Genomics Facility for overseeing the sequencing of our libraries. We thank Chao Zhang (Department of Chemistry, University of Southern California), who originally advised us to make the N84T mutation for the generation of the *mcs6*^*as5*^ strain. We are grateful for valuable discussions with Charles Danko and Hojoong Kwak regarding transcription wave calling and other computational methods.

## Author Contributions

G.T.B. performed PRO-seq experiments and corresponding data analysis. P.P. conducted and analyzed biochemical assays. M.S. created the *mcs6*^*as5*^ strain. G.T.B., P.P., R.P.F, and J.T.L. designed experiments and prepared the manuscript.

## Author Information

Reprints and permissions information is available at http://www.nature.com/reprints. Correspondence and requests for materials should be addressed to

## Methods

### Yeast strains

Yeast strains are listed in Extended Data Table 1.

### Construction of *mcs6*^*as5*^ mutant fission yeast

We previously described *mcs6* and *cdk9* AS mutant strains, each with single substitutions of the gatekeeper residues, Leu87 and Thr120, respectively, with Gly^21^. Although growth of both strains was sensitive to 3-MB-PP1, the *mcs6*^*L87G*^ strain was ~5-fold less sensitive than was the *cdk9*^*T120G*^ strain^21^. By standard methods for gene replacement in *S. pombe*^24^, we introduced structure-guided second site mutations to attempt to optimize performance of *mcs6*^*as*^ alleles^25^ and obtained a double-point mutant, *mcs6*^*N84T/L87G*^ (referred to hereafter as *mcs6*^*as5*^), which had a growth rate indistinguishable from that of wild-type cells in the absence of 3-MB-PP1, and was sensitive to growth inhibition by 3-MB-PP1 with an IC_50_ ≈ 5 μM, which was nearly identical to that of *cdk9*^*as*^ (Extended Data Fig. 3a).

### Monitoring loss of target phosphorylation upon kinase inhibition

For measuring loss of phosphorylation on Spt5 and Rpb1 upon CDK inhibition, cell lysates were rapidly prepared using the trichloroacetic acid (TCA) lysis method^26^. Briefly, cultures were grown to a density of ~1.2 x10^7^ cells/ml in YES media at 30°C prior to treatment. After the specified duration of treatment with of 3-MB-PP1 or DMSO, ~ 6 x 10^7^ cells were transferred into 500 μl 100% (w/v) TCA and collected by centrifugation. Pellets were resuspended in 20% (w/v) TCA and vortexed for the preparation of protein extracts. Antibodies used in this study recognized Spt5-P or total Spt5^27^, total Pol II (BioLegend, MMS-126R), Pol II Ser2-P (Abcam, ab5095), Pol II Ser5-P (BETHYL A304-408A), Pol II Ser7-P (EMD Millipore 04-1570), and α-tubulin (Sigma, T-5168)

### Treatment of analog sensitive strains for PRO-seq

All samples were grown and treated in YES medium at 30 °C. Biological replicates were prepared by picking distinct colonies of a particular strain and growing them in separate liquid cultures. For each experiment, cultures were grown overnight and diluted to an optical density OD_600_ = 0.2. Diluted cultures were grown to mid-log phase (OD_600_ = 0.5-0.6), before treatments. Prior to treatment, all culture volumes were adjusted (based on OD_600_) to have an equal number of *S. pombe* cells (in 10 mL volume). After normalization for cell number, a fixed amount of thawed *S. cerevisiae* (50 μL OD_600_ = 0.68) culture was spiked in to each sample. Treatments were performed by adding either 10 μM 3-MB-PP1 (stock concentration = 40 mM) or an equivalent volume of DMSO. Sample treatments were terminated by pouring into 30 mL ice-cold water, treated with diethyl pyrocarbonate DEPC. Samples were immediately spun down and processed for cell permeabilization as described previously ^4^.

### Time course experiments

The time course experiment was performed in biological replicates, where all time-points for a given replicate was performed on cells from the same large culture. Each time-point treatment was performed as described above. In order to limit differences in time on ice before permeabilization, treatment time-points were carried out in reverse order. The 2.5, 1, and 0.5-minute treatments were performed separately to ensure accuracy of timing. The 0 minute time point was treated with DMSO for 20 minutes to account for any possible effects of adding the solvent for the maximum duration.

### PRO-seq

Precision run-on sequencing libraries in fission yeast strains with spiked-in *S. cerevisiae* for normalization were prepared as described previously^4,28^. For specific batches of experiments slight modifications to the library preparation were made. Samples prepared with *wt, mcs6*^*as5*^, or *cdk9*^*as*^ outside of the time course experiment were prepared according to the standard procedure^28^. All libraries prepared from *lsk1*^*as*^ mutant and *mcs6*^*as5*^ *cdk9*^*as*^ double mutant strains received a novel 3’ RNA adaptor during the first RNA ligation step. This 3’ adaptor contains a random 6 nt molecular barcode at the 5’ end that, although not used here, can be used during read processing to remove PCR duplicates (5’-/5Phos/NNNNNNGAUCGUCGGACUGUAGAACUCUGAAC/Inverted dT/-3’). For all time-course samples prepared in *cdk9*^*as*^, as well as *lsk1*^*as*^ *cdk9*^*as*^ samples, a different 3’ RNA adaptor was used. Here the RNA oligonucleotide possessed a known hexanucleotide sequence preceded by a guanine at the 5’ end, which differed for each library prepared (5’ -/5Phos/G<u>NNNNNN</u>GAUCGUCGGACUGUAGAACUCUGAAC-/Inverted dT/). The ligation of this adaptor with distinct barcodes to each library permitted the pooling of all libraries after the 3’ end ligation step and thus facilitated handling. After pooling, the remainder of the library preparation was carried out as usual, but in a single tube. After sequencing, the inline barcode was used to parse reads based on their sample of origin.

### Alignment and data processing

All sequencing was performed on an Illumina NextSeq 500 device. If samples were pooled during the library preparation using the novel 3’ RNA adaptor described above, raw reads were parsed into their respective samples based on the inline barcode using fastx_barcode_splitter function from the FASTX-toolkit (http://hannonlab.cshl.edu/fastx_toolkit/). Raw reads from each sample were processed by removing any instances of partial or complete matches to the 5’ adaptor sequence (5’-TGGAATTCTCGGGTGCCAAGG -3’) with the fastx_clipper function. Next, if a 3’ molecular, or sample barcode was included during 3’ adaptor ligation, this length was removed from the beginning of each read. All reads were then trimmed to a maximum remaining length of 36 nt using the fastx_trimmer function. With the fastx_reverse_complement function we retrieved the reverse complement sequence of each read. All downstream alignments were performed using Bowtie (version 1.0.0)^29^. Reverse complemented reads derived from ribosomal RNA genes were then removed through alignment to a genome consisting of ribosomal RNA genes from *Saccharomyces cereivisiae* (spike-in) and then *Schizosaccharomyces pombe*. To parse reads based on organism of origin, while eliminating reads of ambiguous origin, non-ribosomal DNA reads were then aligned to a combined genome, containing each chromosome from *S. cerevisiae* (sacCer3 = S288C_reference_genome_R64-1-1_20110203) and *S. pombe* (version: ASM294v2). Only uniquely mapping reads were considered for downstream analysis. Ultimately, the normalization factor for each library was calculated as: total spike-in mapped reads/10^5^. Unique read alignments to the fission yeast genome were processed using the Bedtools^30^ function, genomeCoverageBed, to generate bedgraph formatted files, reporting the number of read 3’ ends (the last base incorporated) at each position across the genome. Bedgraph files were further converted to bigwig format for downstream analysis. Sequencing, alignment, and batch information for each sample can be found in Extended Data Table 2.

### Combined replicates

Biological replicates correlated very well with spike-in normalization centering gene by gene scatter around the line x=y (Extended data Fig. 2), allowing us to combine the raw data from replicates (pre-alignment) for added read depth when useful. In one particular sample, *mcs6*^*as5*^ treated with 3-MB-PP1 for 5 minutes, we ultimately omitted biological replicate one (Extended data Table 2) from all analysis in this work, including composite profiles, due to concerns with the library quality. Importantly, limiting differential expression analysis between treated and untreated samples to only one replicate in the treated condition reduces power to detect differences due to overestimates of variance^13^. Although the number of real changes in the tested gene regions after inhibition of *mcs6*^*as*^ may be higher than listed in Figure 2e, the reported global shifts in Pol II distribution can be seen in individual replicates.

### Experimental batches

Three “batches” of experiments were performed to generate the data in this work (Extended Data Table 2). In order to minimize noise introduced from across-batch comparisons, all analyses presented herein were restricted to PRO-seq libraries prepared within the same batch.

### Gene sets

All genes used in this work were required to have an observed transcription start site, as previously identified (n = 4672)^4^, and this observed TSS was used throughout. We then filtered genes for “activity” in our untreated, wild type data. A gene was considered active if the read density within the gene body was significantly higher (p < 0.01) than a set of intergenic “background” regions, based on a Poisson distribution with λ = background density (λ = 0.0402; n active = 4589). Finally, to minimize the possible influence of read-through transcription from upstream genes, we filtered genes based on the relative amounts of PRO-seq signal directly upstream of the TSS. Thus, we only considered genes with more downstream (+250 to +550) relative to immediately upstream (-300 to observed TSS; n = 3330).

### Differential expression analysis

Raw reads from the appropriate strand were counted within specified windows using custom scripts. Differential expression analysis was performed for desired regions using the DESeq2 R package^13^. Rather than using default between-sample normalization approaches, we supplied our own spike-in based normalization factors (Extended data table 2) using the “sizeFactors” argument. Genes and regions were considered significantly changed (up or down) if they were computed to have an adjusted p-value < 0.01.

### Advancing and clearing wave analysis

Both advancing and clearing waves were identified using a three state Hidden Markov Model (HMM) that was previously developed and implemented on GRO-seq data from a human cell line^18^. To increase the number of time points for which we could identify advancing waves, we restricted our analysis to calling waves on filtered genes longer than 4 kb (n = 224) at each time point up to five minutes. Wave calling parameters were adjusted to accommodate the smaller genes of yeast. For calling advancing waves, the upstream region used was set to -500 to the observed TSS for each gene. Approximate wave distances, used to initialize the model, were set based on estimates derived from inspecting the fold-change heat maps (30 sec = 1 kb; 1 min = 1.5 kb; 2.5 min = 2.5 kb; 5 min = 3.5 kb). Wave distances were determined based on the difference in signal within 50 bp windows between treated and untreated. The wave quality was determined based on the criteria used previously^18^. We further required that for each gene, a wave must be called for all used time points (maximum 5 minutes) using the combined replicate data, and that the calls must not recede toward the TSS with time (n = 43).

To identify regions containing the faster population of Pol II (i.e. the clearing wave), we adjusted the HMM to identify the region between the advancing and clearing waves. For this task we used the first 1000 nt downstream of the observed TSS (within the advancing wave) to initialize the first state of the HMM. Unlike the model used to call the advancing wave, which assumed the upstream state to be normally distributed, all three states were presumed to behave according to distinct gamma distributions. Because of the fleeting nature of this population of polymerases observed in our time course, we were further limited to analysis of filtered genes longer than 6 kb (n = 38). Again, 50 nt windows were used to tile across genes, however an additional smoothing parameter in the model arguments (TSmooth) was set to 5 as a way of restricting the effect of windows with outlying differences between treated and untreated. For each gene, we required that the model be able to identify a distinct clearing wave (Kullback-Leibler divergence > 1, between states 2 and 3) at each time point and that did not move towards the TSS with time (n = 28).

We employed two methods to assess the confidence in measured average rates. For the first, bootstrapping approach, an average rate was calculated using 10% of genes for which we called waves. This process was repeated 1000 times (with replacement), giving a distribution of means. A more conservative, gene-by-gene estimate of variance was derived simply from the distribution of rates based on individual measurements for each gene.

## Data Access

The raw and processed sequencing files have been submitted to the NCBI Gene Expression Omnibus (GEO; http://www.ncbi.nlm.nih.gov/geo/) under accession GSE102308

## Code Availability

Custom scripts and alignment pipelines have been made publicly available through the following GitHub repository; https://github.com/gregtbooth/Pombe_PROseq.

**Extended Data Figure 1.**
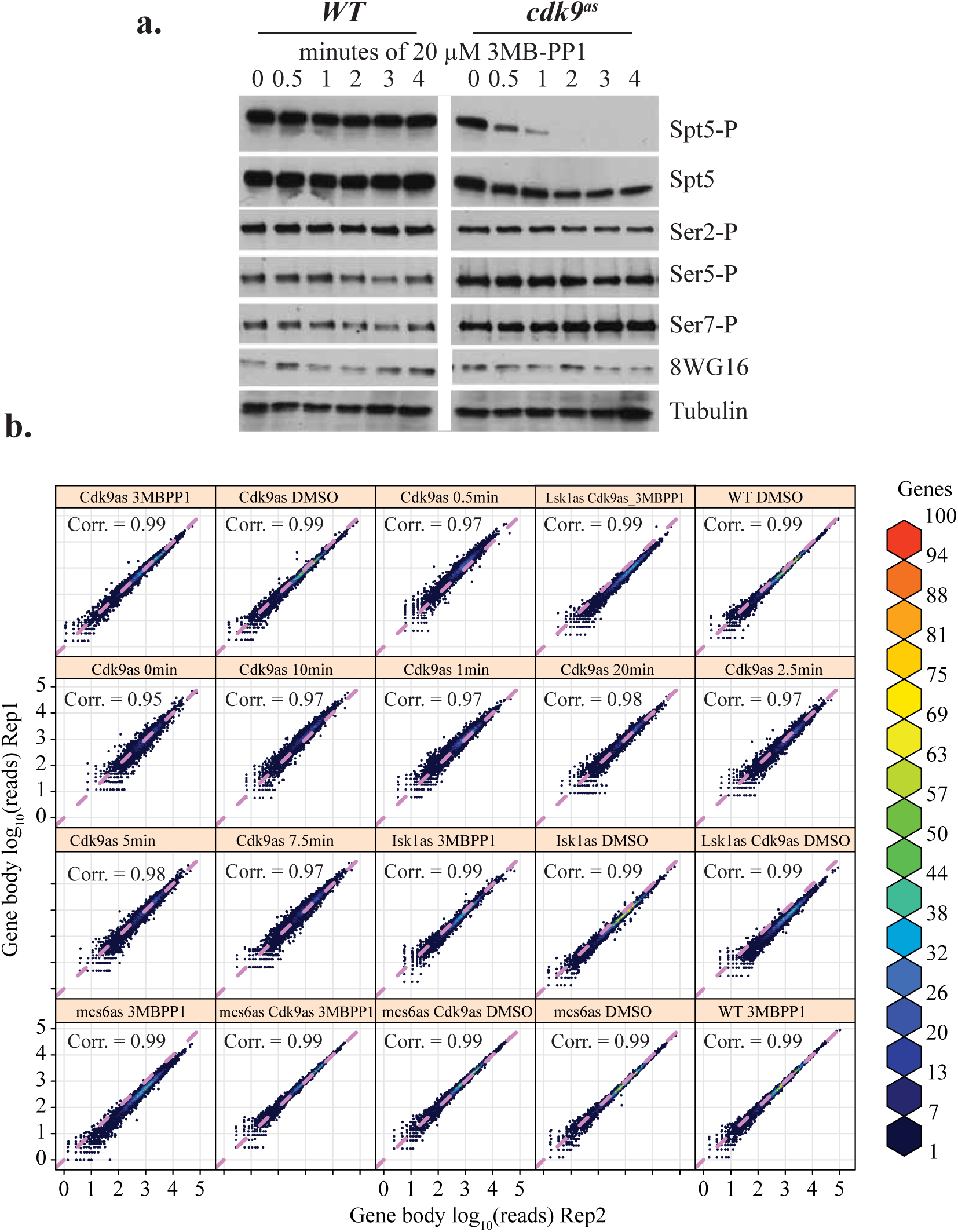
Cdk9 inhibition primarily affects Spt5-P within a short timeframe and PRO-seq experiments are reproducible. **a.** Western blot analysis of phosphorylated residues within the elongation complex in *wt* and *cdk9*^*as*^ strains after increasing durations of treatment with 20 μM 3-MB-PP1. From top to bottom, antibodies were raised against, Spt5-P, total Spt5, Ser2-P, Ser5-P, Ser7-P, Rpb1, or Tubulin (loading control). **b.** Scatter plots displaying a correlation (Spearman’s rho) between biological replicate PRO-seq data for each sample. Data points show spike-in normalized read counts (log_10_) within the gene body (TSS + 200 bp to annotated CPS) of all filtered genes. Since minimal variation is expected between replicates, accurate spike in-based normalization will produce scatter that is approximately centered on the purple diagonal line (x = y).

**Extended Data Figure 2.**
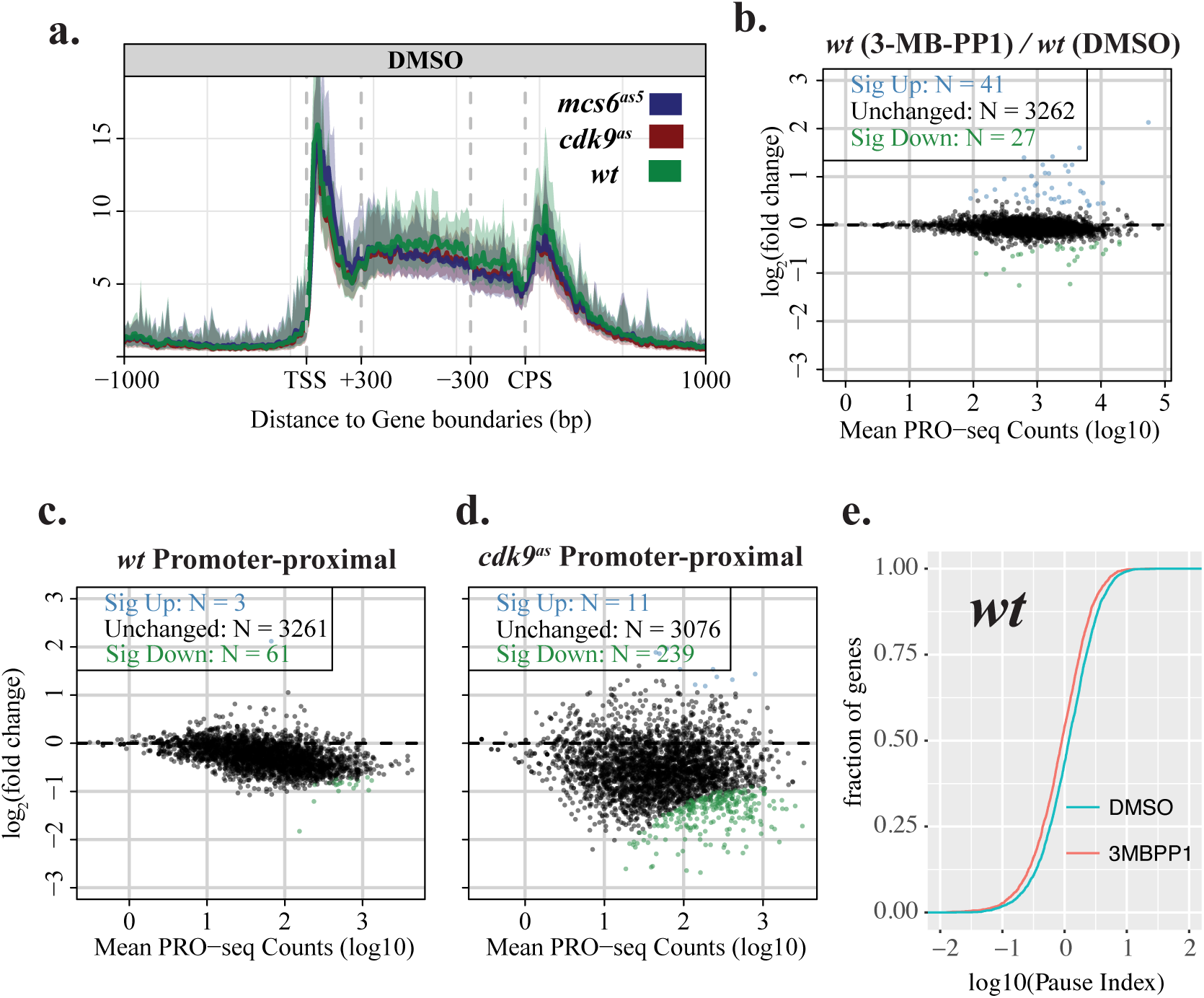
AS kinase-dependent strains provide a highly controlled system for studying transcription. **a.** Composite profiles of combined replicate data for samples of untreated wild type, *mcs6*^*as5*^, and *cdk9*^*as*^ strains across filtered genes separated from nearest same strand neighbors by at least 1 kb on both sides (n = 919). Composite profiles represent the median subsampled value within each bin. Shaded regions correspond to the 12.5% and 87.5% quantiles. To scale genes to a common length, the middle gene body region of each gene was scaled to 60 bins, while the regions -1000 bp to +300 bp relative to the TSS, and -300 bp to +1000 bp relative to the CPS, are unscaled 10 bp windows. **b.** MA plot displaying the log_2_ fold change between gene body read counts from treated and untreated *wt* cells, as calculated using the DESeq2 package. **c & d.** MA plot displaying the log_2_ fold change between treated and untreated promoter proximal (TSS to +100 bp) read counts for wild type (**c.**), or *cdk9*^*as*^ (**d.**). Genes that show significantly increased or decreased PRO-seq signal (adjusted *p* <0.01) are shown in blue and green, respectively. For all MA plots, spike-in normalization was used when calculating log_2_ fold changes between samples. **e.** Cumulative density functions for pausing index (log_10_) of all filtered genes in treated (red) and untreated (blue) samples for the wild type strain.

**Extended Data Figure 3.**
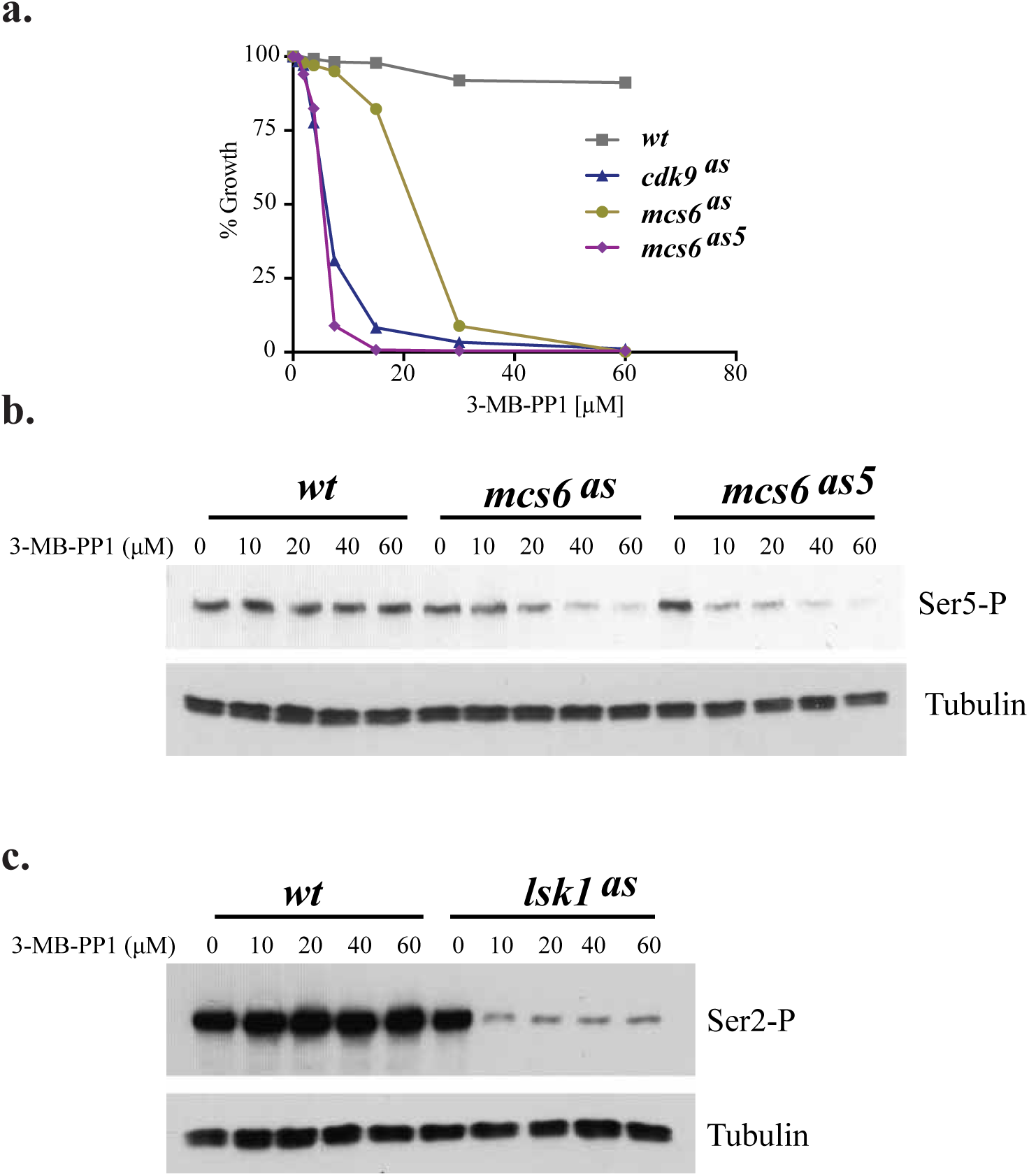
Characterization of AS strains and the effects of 3-MB-PP1 concentration. **a.** The indicated strains were grown in 96-well plates in the presence of increasing amounts of 3-MB-PP1 and their growth quantified by OD_600_. The mutant alleles and their associated mutations are: *mcs6*^*as*^, L87G; *mcs6*^*as5*^, N84T/L87G; *cdk9*^*as*^, T120G. **b.** Western blot analysis of Ser5-P relative to tubulin (loading control) in *wt*, *mcs6*^*as*^, and *mcs6*^*as5*^ strains after 1 hr treatment at 30 °C with the specified concentration of 3-MB-PP1. **c.** Western blot analysis of Ser2-P relative to tubulin in *wt*, and *lsk1*^*as*^ strains after 1 hr of treatment at 30 °C with the specified concentration of 3-MB-PP1.

**Extended Data Figure 4.**
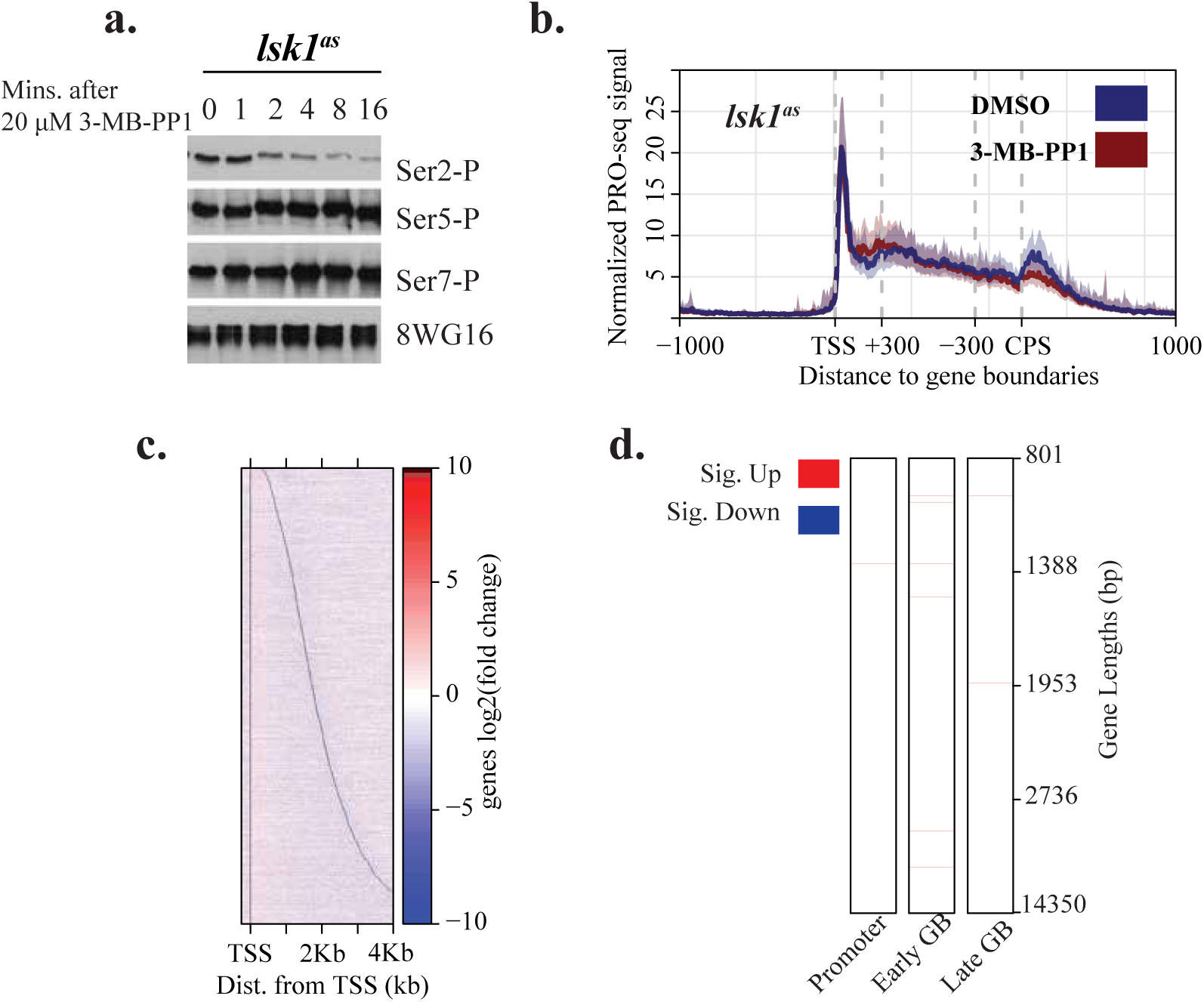
Treatment of *lsk1*^*as*^ with 3-MB-PP1 has minimal impact on transcription within five minutes, despite loss of Ser2-P. **a.** Western blot analysis of phosphorylated CTD residues (Ser2-P, Ser-5-P, Ser7-P) in relation to total CTD signal. Levels were measured over a time course of *lsk1*^*as*^ treated with 20 μM 3-MB-PP1. **b.** Composite PRO-seq profiles for treated (5 min 10 μM 3-MB-PP1) and untreated (5 min DMSO) *lsk1*^*as*^ displaying median subsampled signal across filtered genes separated from nearest same strand neighbors by at least 1 kb on both sides (n = 919). Shaded regions correspond to the 12.5% and 87.5% quantiles. The middle gene body region of each gene was scaled to 60 bins, while the regions, -1000 bp to +300 bp, relative to the TSS, and -300 bp to +1000 bp relative to the CPS, are unscaled 10 bp windows. **c.** Heat map depicting the log_2_ fold change in normalized PRO-seq signal (treated/untreated) in *lsk1*^*as*^ within 10 bp bins from -250 bp to +4000 bp relative to the TSS. Genes are sorted by increasing length, with black lines representing observed TSS and annotated CPS. **d.** Heat maps depicting whether each gene exhibits a significant fold change (adjusted *p* < 0.01; treated/untreated) in promoter, early, or late gene body regions (defined in Figure 2). Genes were required to be longer than 800 nt and are sorted from top to bottom by increasing gene length. Gene length quartiles are shown with tick marks on the right. Significant increases and decreases in each region are shown as red and blue, respectively.

**Extended Data Figure 5.**
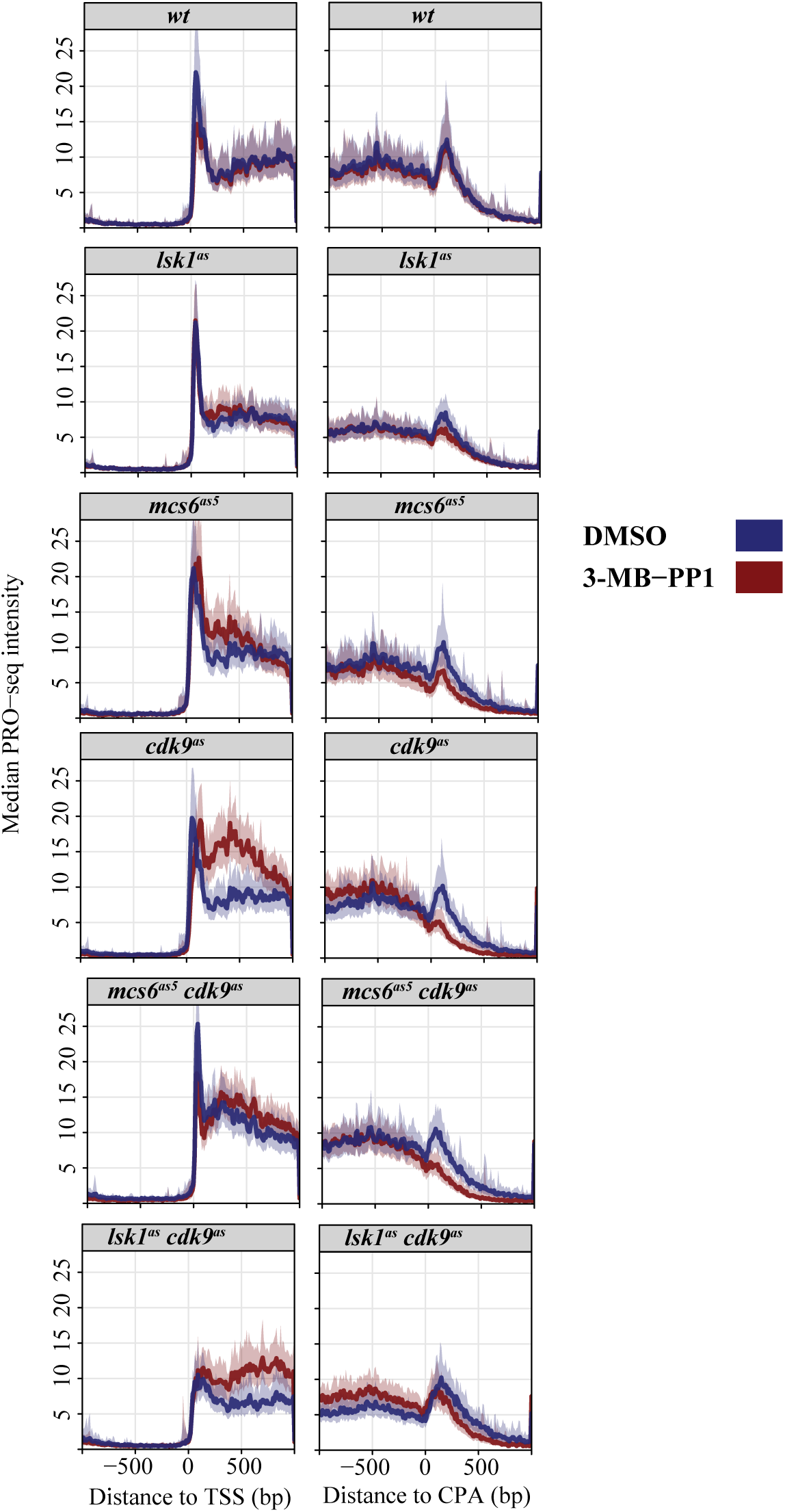
Global effects of kinase inhibition revealed through composite profiles centered on TSS and CPS. Each panel displays centered composite PRO-seq profiles of treated (5 min 10 μM 3-MB-PP1) and untreated (5 min DMSO) samples from each strain. Dark middle lines reflect the median sub-sampled PRO-seq signal within 10 bp windows from -1 kb to +1 kb from observed TSS (left) or CPS (right) for genes separated from nearest same strand neighbors by at least 1 kb on both sides (n = 919). Shaded regions correspond to the 12.5% and 87.5% quantiles.

**Extended Data Figure 6.**
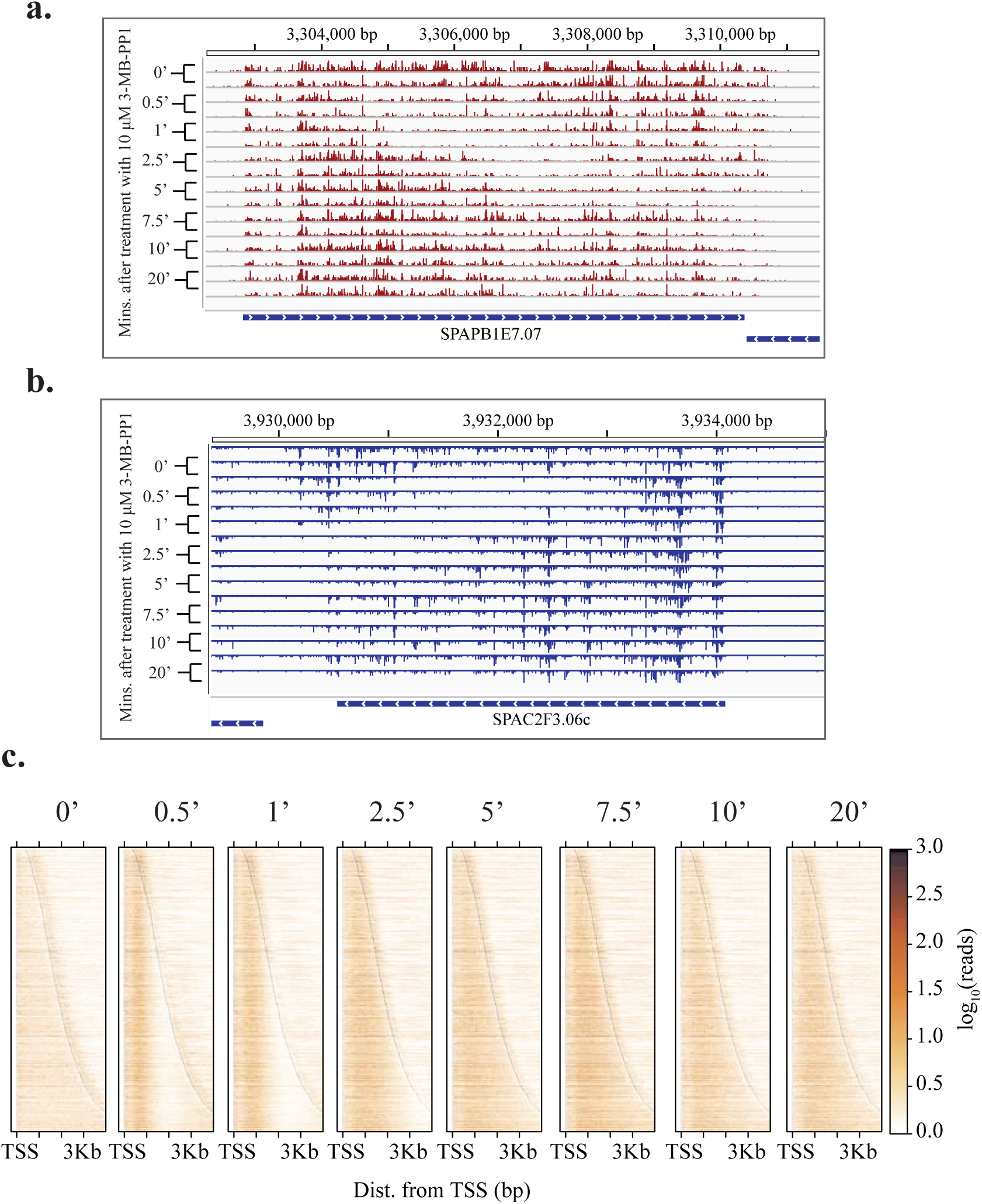
Reproducible time-dependent impact of Cdk9 inhibition on transcription. **a & b.** Browser track images from the *SPAPB1E7.07* (**a.**) and *SPAC2F3.06c* (**b.**) loci. Tracks represent normalized read counts from biological replicates of *cdk9*^*as*^ on the plus or minus strands, respectively, treated with 10 μM 3-MB-PP1 for increasing amounts of time. **c.** Raw heatmaps of PRO-seq signal for strains before and after treatment. Heatmap signal intensity reflects normalized signal (log_10_) within 10 bp bins from -250 to +4000 bp relative to the TSS for data from combined replicates for increasing treatment durations in *cdk9*^*as*^, from left to right. Within each panel, genes are sorted by increasing length from top to bottom, with dark lines showing boundaries for each gene. All samples were treated with 10 μM 3-MB-PP1 for the specified times. The zero minute treatment was treated with DMSO for 20 min.

**Extended Data Figure 7.**
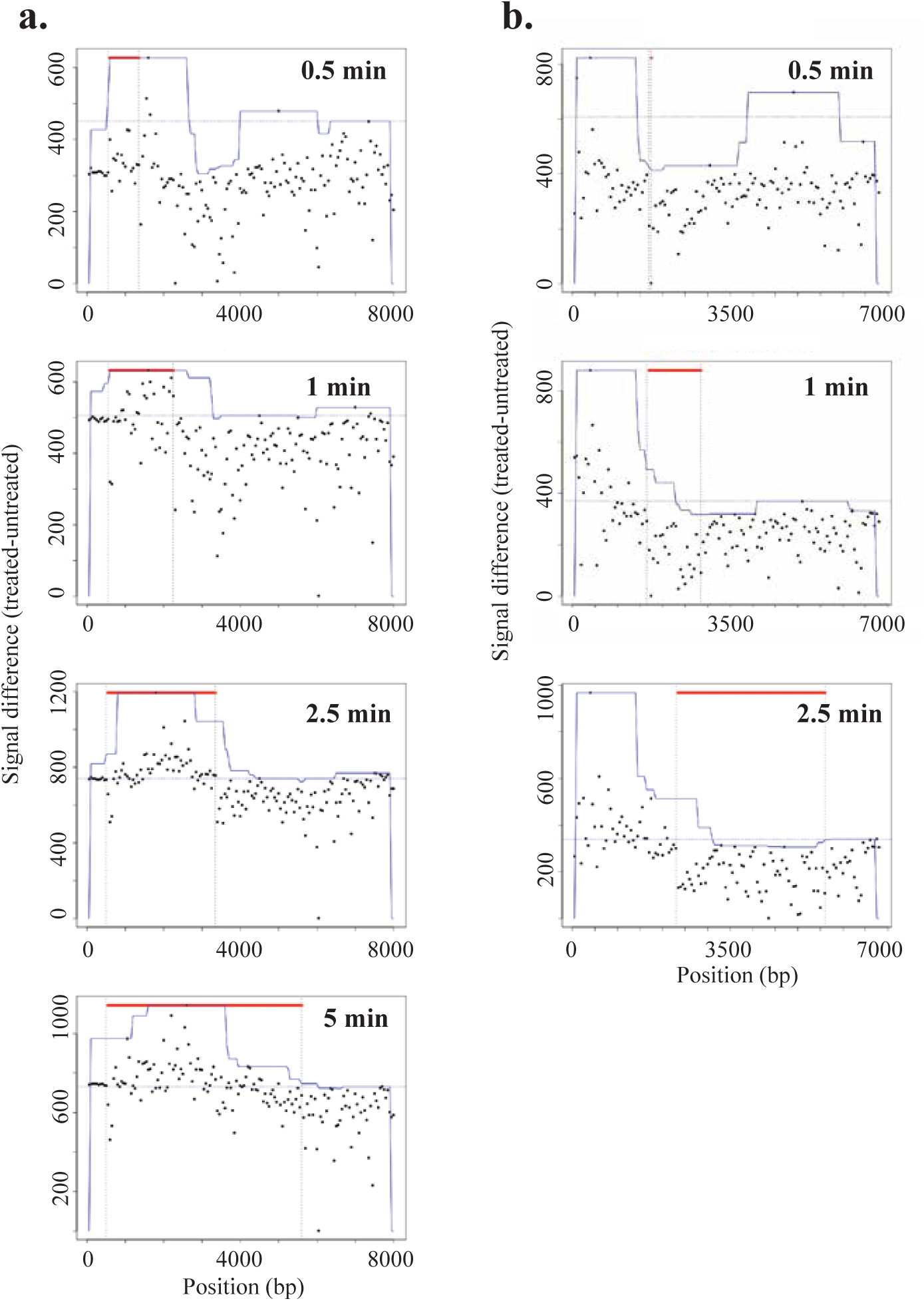
Examples of difference maps used for calling waves at each time point. **a.** Advancing wave location estimates based on difference in PRO-seq signal (treated – untreated) measured within 50 bp windows over the *SPAPB1E7.07* locus at each time point (0.5 min, 1 min, 2.5 min, and 5 min). **b.** Estimates of region between advancing and clearing waves based on difference in PRO-seq signal (treated – untreated) measured within 50 bp windows over the *SPAC926.09c* locus at each time point (0.5 min, 1 min, 2.5 min). For each plot, the y-axis displays the difference within each 50 bp window plus a constant (dotted horizontal line), which makes the minimum value equal to one. The red points indicate windows identified as being within the advancing wave (a) or between waves (b). The blue trace reflects the local maximum within a moving 1 kb window.

**Extended Data Figure 8.**
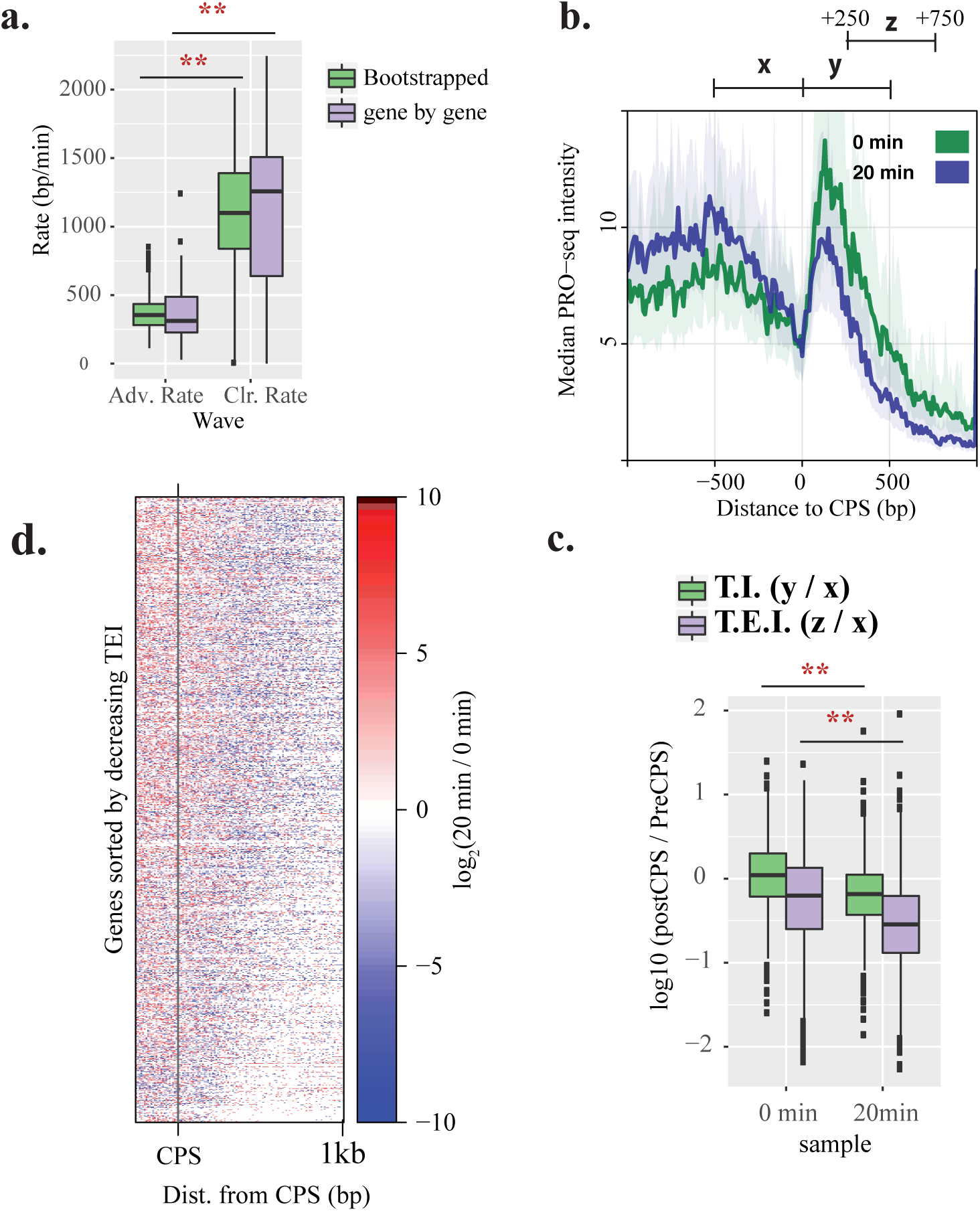
Decreased Pol II transcription rates globally reduce post-CPS elongation upon continued Cdk9 inhibition. **a.** Box plots show distributions of rate estimates for the advancing wave (left) and clearing wave (right). Significant differences were observed between advancing and clearing rates using both bootstrapping (*p* < 2.2e-16) and gene-by-gene (*p* = 7.719e-6) approaches (Student’s t-test). **b.** CPS centered composite PRO-seq profiles comparing *cdk9*^*as*^, either untreated (green) or after 20 min of treatment with 10 μM 3-MB-PP1 (blue). Dark middle lines reflect the median subsampled PRO-seq signal within 10 bp windows from -1 kb to +1 kb of the CPS for genes separated from nearest same strand neighbors by at least 1 kb on both sides (n = 919). Shaded regions correspond to the 12.5% and 87.5% quantiles. **c.** Distributions of termination index (T.I.; green) and termination elongation index (T.E.I., purple) for all genes used in (a) before (left) and after 20 min of Cdk9 inhibition (right). T.I. and T.E.I. are metrics that capture elongation beyond the CPS, relative to signal in the late gene body. All 500 bp regions used for calculating T.I. (y / x) and T.E.I. (z / x) ratios are depicted above the plot in (b). **d.** Heatmap of log_2_ fold change (20 min/ 0 min) within 10 bp windows from -250 to +1000 bp relative to CPS for genes used in (b) and (c). Genes are sorted by decreasing T.E.I. from top to bottom as calculated from untreated *cdk9*^*as*^. Data from combined replicates was used for all plots and calculations.

**Extended Data Table 1.** Yeast strains

| Strain | Alias | Genotype | Reference |
| --- | --- | --- | --- |
| JS78 | *wt* | *leu1-32 ura4-D18 his3-D1 ade6-M210 h<sup>+</sup>* | Saiz & Fisher 2002 |
| LV7 | *cdk9<sup>as</sup>* | *cdk9<sup>T120G</sup>::kanMX6 leu1-32 ura4-D18 his3-D1 ade6-M210 h<sup>+</sup>* | Viladevall et al. 2009 |
| MS340 | *mcs6<sup>as5</sup>* | *mcs6<sup>N84T-L87G</sup>::kanMX6 leu1-32 ura4-D18 his3-D1 ade6-M210 h<sup>+</sup>* | This study |
| CS118 | *lsk1<sup>as</sup>* | *lsk1<sup>F353G</sup>::kanMX6 leu1-32 ura4-D18 his3-D1 ade6-M216 h<sup>-</sup>* | Viladevall et al. 2009 |
| PP20 | *cdk9<sup>as</sup> lsk1<sup>as</sup>* | *cdk9<sup>T120G</sup>::hphMX6 lsk1<sup>F353G</sup>::kanMX6 leu1-32 ura4-D18 his3-D1 ade6-M210 h<sup>+</sup>* | This study |
| PP23 | *cdk9<sup>as</sup> mcs6<sup>as5</sup>* | *cdk9<sup>T120G</sup>::hphMX6 mcs6<sup>N84T-L87G</sup>::kanMX6 leu1-32 ura4-D18 his3-D1 ade6-M210 h<sup>-</sup>* | This study |
| *S. cerevisiae* | Spike-in | W303-1a | Booth et al. 2016 |

**Extended Data Table 2.** PRO-seq sample information

| sample | spike-in<br>ribosomal | ribosomal | spike-in | genome | total | Norm.<br>Factor | batch |
| --- | --- | --- | --- | --- | --- | --- | --- |
| WT_DMSOr1 | 10,174,782 | 12,285,660 | 46,236 | 6,157,670 | 30,465,302 | 0.46236 | 1 |
| WT_DMSOr2 | 12,204,603 | 15,627,984 | 54,480 | 8,401,775 | 38,708,110 | 0.5448 | 1 |
| WT_3MBPP1_1 | 12,326,132 | 15,585,548 | 80,667 | 10,653,445 | 41,242,195 | 0.80667 | 1 |
| WT_3MBPP1_2 | 10,582,750 | 13,617,874 | 62,899 | 8,593,076 | 35,067,313 | 0.62899 | 1 |
| mcs6_DMSO1 | 12,257,633 | 15,672,640 | 80,616 | 10,205,273 | 41,454,045 | 0.80616 | 1 |
| mcs6_DMSO2 | 11,707,207 | 14,554,011 | 56,817 | 8,051,885 | 36,735,682 | 0.56817 | 1 |
| mcs6_3MBPP1_1* | 11,430,584 | 13,874,952 | 101,502 | 6,331,691 | 37,568,528 | 1.01502 | 1 |
| mcs6_3MBPP1_2 | 12,052,555 | 14,402,612 | 72,480 | 8,466,890 | 37,221,280 | 0.7248 | 1 |
| CDK9_DMSO1 | 12,389,079 | 14,034,425 | 54,917 | 6,367,375 | 34,927,058 | 0.54917 | 1 |
| CDK9_DMSO2 | 12,778,993 | 16,323,383 | 66,519 | 9,358,197 | 41,353,473 | 0.66519 | 1 |
| CDK9_3MBPP1_1 | 11,143,318 | 13,228,901 | 60,777 | 7,827,810 | 34,514,302 | 0.60777 | 1 |
| CDK9_3MBPP1_2 | 12,517,108 | 16,637,590 | 88,178 | 12,311,216 | 45,190,617 | 0.88178 | 1 |
| Isk1as_DMSOr1 | 8,137,437 | 9,062,002 | 73,907 | 6,843,886 | 29,144,786 | 0.73907 | 2 |
| Isk1as_DMSOr2 | 10,483,655 | 11,431,617 | 67,401 | 8,289,801 | 35,769,690 | 0.67401 | 2 |
| Isk1as_3MBPP1r1 | 6,935,506 | 7,256,537 | 54,000 | 5,481,567 | 23,207,739 | 0.54 | 2 |
| Isk1as_3MBPP1r2 | 3,080,010 | 2,899,634 | 17,547 | 2,063,159 | 12,330,930 | 0.17547 | 2 |
| Mcs6as_CDK9as_DMSOr1 | 7,733,612 | 8,014,461 | 37,501 | 5,750,563 | 26,364,431 | 0.37501 | 2 |
| Mcs6as_CDK9as_DMSOr2 | 8,149,246 | 8,775,629 | 41,878 | 5,992,027 | 26,861,563 | 0.41878 | 2 |
| Mcs6as_CDK9as_3MBPP1r1 | 7,206,380 | 7,247,941 | 33,500 | 4,536,765 | 23,488,806 | 0.335 | 2 |
| Mcs6as_CDK9as_3MBPP1r2 | 6,378,728 | 6,817,084 | 36,155 | 4,582,814 | 20,834,265 | 0.36155 | 2 |
| Cdk9as_0min_DMSO_rep1 | 941,848 | 1,294,433 | 8,443 | 1,240,837 | 6,521,490 | 0.08443 | 3 |
| Cdk9as_0min_DMSO_rep2 | 1,065,358 | 1,504,048 | 13,568 | 2,005,311 | 10,353,030 | 0.13568 | 3 |
| Cdk9as_0.5min_10uM_3MBPP1_rep1 | 3,205,493 | 3,805,818 | 24,773 | 3,807,026 | 15,454,733 | 0.24773 | 3 |
| Cdk9as_0.5min_10uM_3MBPP1_rep2 | 1,139,637 | 1,662,171 | 16,380 | 1,738,590 | 7,779,488 | 0.1638 | 3 |
| Cdk9as_1min_10uM_3MBPP1_rep1 | 1,699,913 | 2,207,482 | 16,573 | 2,128,619 | 9,584,199 | 0.16573 | 3 |
| Cdk9as_1min_10uM_3MBPP1_rep2 | 2,130,628 | 3,151,162 | 30,364 | 3,413,440 | 12,760,335 | 0.30364 | 3 |
| Cdk9as_2.5min_10uM_3MBPP1_rep1 | 1,753,947 | 2,386,420 | 14,135 | 1,973,114 | 8,645,482 | 0.14135 | 3 |
| Cdk9as_2.5min_10uM_3MBPP1_rep2 | 2,557,168 | 3,377,095 | 23,249 | 3,621,442 | 14,120,294 | 0.23249 | 3 |
| Cdk9as_5min_10uM_3MBPP1_rep1 | 2,394,714 | 3,307,974 | 20,338 | 2,812,736 | 15,961,497 | 0.20338 | 3 |
| Cdk9as_5min_10uM_3MBPP1_rep2 | 2,504,115 | 2,672,961 | 26,141 | 3,742,474 | 17,870,169 | 0.26141 | 3 |
| Cdk9as_7.5min_10uM_3MBPP1_rep1 | 1,998,776 | 2,768,515 | 15,073 | 2,612,802 | 16,268,203 | 0.15073 | 3 |
| Cdk9as_7.5min_10uM_3MBPP1_rep2 | 2,662,662 | 3,131,133 | 33,314 | 5,316,423 | 21,503,832 | 0.33314 | 3 |
| Cdk9as_10min_10uM_3MBPP1_rep1 | 2,228,285 | 2,681,515 | 17,655 | 3,070,909 | 15,621,102 | 0.17655 | 3 |
| Cdk9as_10min_10uM_3MBPP1_rep2 | 1,000,575 | 1,441,581 | 14,946 | 1,865,992 | 9,003,041 | 0.14946 | 3 |
| Cdk9as_20min_10uM_3MBPP1_rep1 | 1,059,830 | 1,275,738 | 11,469 | 1,802,836 | 8,859,279 | 0.11469 | 3 |
| Cdk9as_20min_10uM_3MBPP1_rep2 | 2,100,685 | 2,635,181 | 22,006 | 3,327,234 | 15,608,295 | 0.22006 | 3 |
| Lsk1as_CDK9as_DMSOr1 | 1,929,149 | 2,510,779 | 32,183 | 2,953,437 | 10,834,859 | 0.32183 | 3 |
| Lsk1as_CDK9as_DMSOr2 | 2,795,832 | 3,489,519 | 34,863 | 4,400,322 | 16,293,462 | 0.34863 | 3 |
| Lsk1as_CDK9as_3MBPP1r1 | 1,566,015 | 2,350,911 | 24,970 | 2,915,731 | 11,162,211 | 0.2497 | 3 |
| Lsk1as_CDK9as_3MBPP1r2 | 2,789,279 | 3,492,754 | 30,235 | 4,278,326 | 18,766,044 | 0.30235 | 3 |

\* Sample omitted from analysis

